# STAR/STARD1: a mitochondrial intermembrane space cholesterol shuttle degraded through mitophagy

**DOI:** 10.1101/2025.04.03.647084

**Authors:** Prasanthi P. Koganti, Amy H. Zhao, Michael C. Kern, Auriole C. R. Fassinou, Vimal Selvaraj

## Abstract

The import of cholesterol to the inner mitochondrial membrane by the steroidogenic acute regulatory protein (STAR/STARD1) is essential for de novo steroid hormone biosynthesis and the acidic pathway of bile acid synthesis. This robust system, evolved to start and stop colossal cholesterol movement, ensures pulsatile yet swift mitochondrial steroid metabolism in cells. Nonetheless, the proposed mechanism and components involved in this process has remained a topic of ongoing debate. In this study, we elucidate the mitochondrial import machinery and structural aspects of STAR, revealing its role as an intermembrane space cholesterol shuttle that subsequently undergoes rapid degradation by mitophagy. This newfound mechanism illuminates a fundamental process in cell biology and provides precise interpretations for the full range of human STAR mutation-driven lipoid congenital adrenal hyperplasia in patients.

**One Sentence Summary:** STAR activates mitochondrial steroid metabolism as a cholesterol shuttle in the intermembrane space and is destroyed by mitophagy.

## INTRODUCTION

The steroidogenic acute regulatory protein (STAR/STARD1) was first identified in 1994 as a protein rapidly synthesized and degraded during steroidogenic responses (*1*). Since then, its fundamental role in facilitating mitochondrial cholesterol import required for steroid hormone biosynthesis has been well demonstrated both *in vitro* (*2*) and *in vivo* (*3*). Despite clear evidence of its biological importance, the precise molecular mechanism through which STAR achieves cholesterol import has remained uncertain, prompting extensive investigation (*4*).

In a landmark study published in 2002, it was proposed that STAR functions externally to mitochondria (*5*), exerting its biological activity from outside the organelle, with eventual mitochondrial matrix targeting only serving as a mechanism to terminate its function (*6*, *7*). This conceptualization subsequently introduced the idea that an additional protein might be necessary to mediate cholesterol transport into mitochondria, and the mitochondrial translocator protein (TSPO), located on the outer mitochondrial membrane, became the primary candidate (*8–10*).

However, our earlier work in 2014 utilizing TSPO knockout mouse models and cells conclusively demonstrated that TSPO is dispensable for mitochondrial cholesterol import during steroidogenesis (*11–13*). This finding, subsequently replicated by independent groups (*14*, *15*), significantly challenged the established view and revitalized an alternative hypothesis – initially proposed by earlier biochemical studies – that STAR functions as a molten globule, transiently interacting with the outer mitochondrial membrane to facilitate cholesterol import (*16*). Recent research has further embraced and built upon this molten-globule model (*17*, *18*). As a result, authoritative reviews and current textbooks predominantly describe STAR as a cytoplasmic protein transiently tethered to the outer mitochondrial membrane (*19*, *20*).

In humans, impaired STAR function arising from various genetic mutations leads to disruptions in steroidogenesis, manifesting clinically as hypomorphic or lethal forms of lipoid congenital adrenal hyperplasia (lipoid CAH) (*21*). Carrier frequencies for common founder mutations in STAR are notably high in Japanese, Korean, and Palestinian populations; one estimate places the frequency of the Q258Stop mutant allele in Japanese individuals at approximately 1 in 200 (*22*). However, biochemical analyses aimed at clarifying how these mutations compromise STAR activity have provided limited insights, and their outcomes are difficult to reconcile with the proposed molten-globule model. Consequently, direct structure-function relationships of human STAR mutations remain poorly understood. The conserved START (STAR-related lipid transfer) domain, crucial for lipid-binding and transfer activity, has been the subject of significant study (*23*). Yet, important regulatory functions embedded within the N-terminal and C-terminal regions of STAR remain unresolved. While an N-terminal mitochondrial targeting sequence (MTS) has been recognized, it was long considered unnecessary for STAR function (*6*, *7*). Similarly, the potential role of STAR’s C-terminal region has typically been discussed only concerning its proposed interactions with the mitochondrial outer membrane (*6*, *7*).

Upon reviewing the literature, we reasoned that some of the challenges encountered in understanding STAR function could stem from experimental models that inadequately represent physiological conditions. Specifically, most studies in this domain have utilized synthetic systems involving non-steroidogenic COS-1 cells co-transfected with STAR and an engineered protein fusion known as F2 (comprising CYP11A1, adrenodoxin, and adrenodoxin reductase) to artificially enable steroidogenesis [first reported in (*24*)]. This F2 system subsequently became a widely adopted experimental approach for evaluating STAR-dependent mitochondrial cholesterol import (*2*)(*3*, *7*, *8*, *22*), despite concerns about its fidelity and whether steroid biosynthesis in this artificial system accurately reflects physiological processes. To overcome these limitations, we developed a physiologically relevant steroidogenic cell model through CRISPR/Cas9-mediated deletion of endogenous STAR in Leydig cells. Leveraging this refined cellular system, we performed targeted reconstitution with STAR mutants and recombinants, enabling precise evaluation of the functionality and intracellular localization of STAR, ultimately defining its exact site of biological activity.

## RESULTS

### A high-fidelity STAR-deleted steroidogenic test system

We first confirmed that STAR targets mitochondria and determined that fusing peptides to the C-terminus does not alter its mitochondrial localization. The full-length (native) STAR- Emerald fusion protein shows thorough colocalization with mitochondria labeled with TOM20- mRuby fluorescence in MA-10 Leydig cells (Fig 1A). Notably, this mitochondrial targeting contrasts with a previous report suggesting that adding even a short C-terminal His-tag to STAR caused cytoplasmic retention in non-steroidogenic COS-1 cells (*6*). To better reflect physiological conditions and reduce experimental artifacts, we developed a steroidogenic MA-10 Leydig cell model for studying STAR function. We utilized CRISPR/Cas9 to specifically target Exon 2, successfully generating STAR knockout (MA-10*^STKO^*) cell lines (Fig S1A). These MA- 10*^STKO^* clones showed a complete absence of STAR protein and fully recapitulated known STAR knockout phenotypes, notably failing to synthesize steroids upon stimulation (Fig 1B). Aside from lacking STAR-dependent steroidogenesis, MA-10*^STKO^* cells retained full steroidogenic capacity upon supplementation with aqueous-soluble 22R-OH-cholesterol (Fig S1B), consistent with the known STAR-bypass mechanism (*25*). Ultrastructural examination of mitochondria in MA-10*^STKO^* cells showed no abnormal morphologies (Fig 1C), an anticipated outcome since STAR is only transiently translated during steroidogenic stimulation in MA-10 cells.

**Fig. 1.**
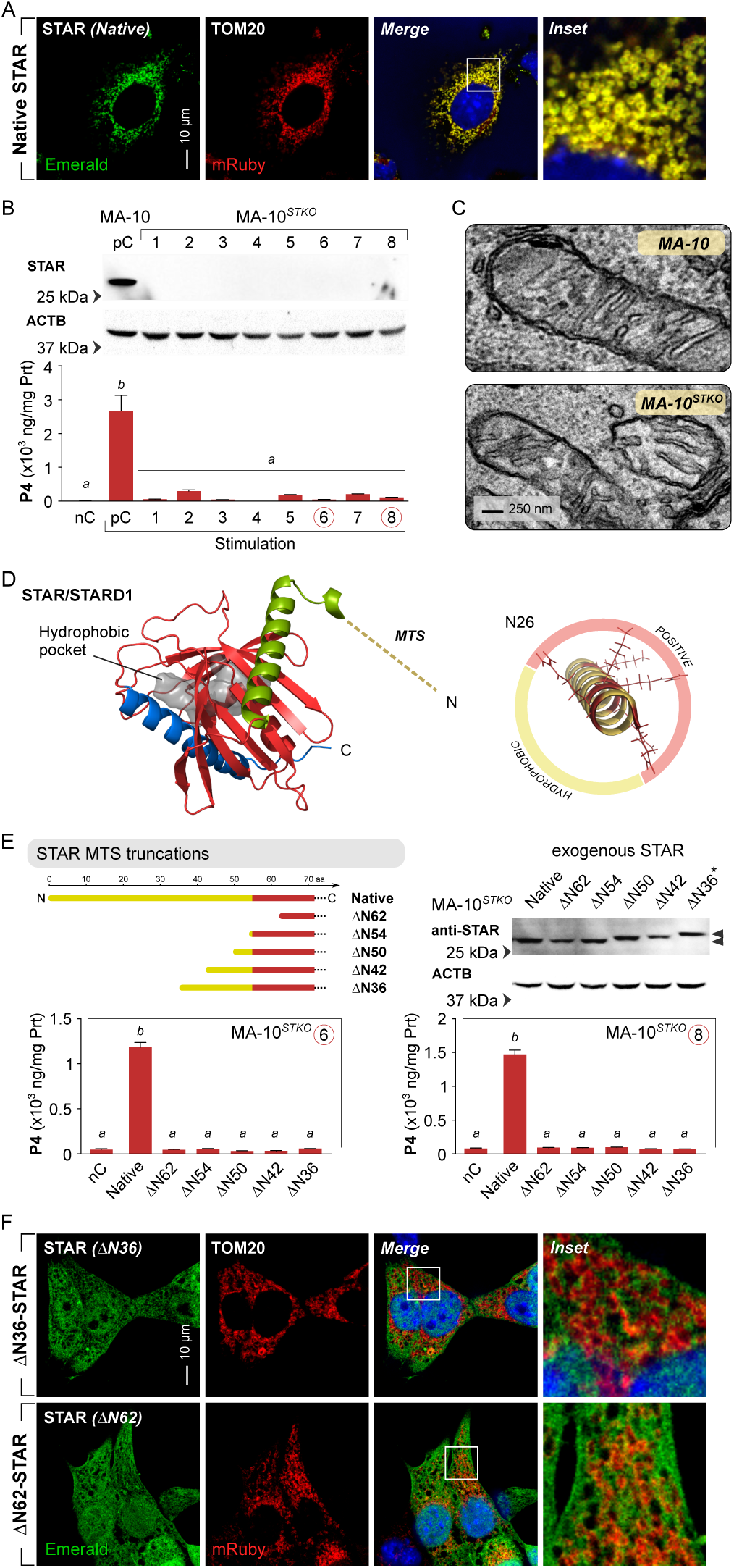
The STAR^MTS^ is essential for mitochondrial cholesterol import function. **(A)** STAR localizes to mitochondria. Confocal imaging shows lentiviral expression of a C-terminal STAR- Emerald fusion protein colocalizes with outer mitochondrial membrane (OMM)-anchored TOM20-mRuby in MA-10 Leydig cells. **(B)** CRISPR/Cas9 deletion of endogenous STAR in MA-10 cells (MA-10*^STKO^*) results in steroidogenic failure after Bt2cAMP stimulation, compared to positive controls (pC) expressing native STAR (also see Fig. S1; negative control, nC; different alphabets indicate significant difference p<0.05). Representative clones (6 and 8) used for subsequent experiments are circled. **(C)** Transmission electron microscopy reveals normal mitochondrial ultrastructure in STAR-deficient MA-10*^STKO^*cells, indicating that STAR deletion does not disrupt mitochondrial morphology. **(D)** Structural model of murine STAR shows an N- terminal α-helix (green) that includes a mitochondrial targeting sequence (MTS), a central hydrophobic cholesterol-binding scaffold of nine anti-parallel β-strands that form the START domain (red), and a C-terminal α-helix with a flexible tail (blue). **(E)** Reconstitution with N- terminal truncation mutants (^ΔN36-^, ^ΔN42-^, ^ΔN50-^, ^ΔN54-^, and ^ΔN62-^STAR) fails to restore steroidogenesis in MA-10*^STKO^*cells compared to native STAR in both clones 6 and 8 (different alphabets indicate significant difference p<0.05). Western blot analysis shows absence of mitochondrial processing as native STAR, particularly evident in ^ΔN36-^STAR, which retains the highest molecular weight (*). **(F)** Confocal microscopy shows that Emerald-tagged truncated STAR proteins (^ΔN36-^STAR and ^ΔN62-^STAR) do not localize to mitochondria, in contrast to native STAR-Emerald [in (A)], confirming that the STAR^MTS^ is essential for its mitochondrial import.

### STAR*^MTS^* is crucial for mitochondrial cholesterol import

Structural examination of murine STAR highlighted several key features (Fig 1D): (a) an N-terminal amphipathic helix characteristic of mitochondrial targeting signals (MTS), (b) a hydrophobic lipid-binding pocket formed by anti-parallel β-sheets and surrounding α-helices, and (c) a flexible, largely unmodeled C-terminal tail. Based on these structural insights, evaluated the functional importance of the N-terminal mitochondrial targeting sequence (STAR*^MTS^*) for STAR localization and activity.

To directly assess the significance of the STAR*^MTS^*, we generated a series of STAR mutants featuring sequential truncations at the N-terminus (^ΔN36-^, ^ΔN42-^, ^ΔN50-^, ^ΔN54-^, and ^ΔN62-^STAR).

Functional analysis in MA-10*^STKO^* Leydig cells demonstrated that none of these truncated STAR constructs could restore stimulated steroidogenesis (Fig 1E). Notably, even the minimal truncation (^ΔN36-^STAR) was ineffective, underscoring the critical importance of the N-terminal sequence for STAR function. These findings stand in marked contrast to previous studies utilizing the F2 system in COS-1 cells, where STAR lacking the entire MTS (^ΔN62-^STAR) retained full steroidogenic capability (*5*, *7*).

Immunoblot analysis revealed that smaller truncations yielded STAR proteins with higher molecular weights (Fig 1E), indicating that the expected post-translational processing of the STAR*^MTS^*, integral for mitochondrial localization/entry, did not occur in these truncated constructs. Importantly, the complete STAR*^MTS^* deletion (^ΔN62-^STAR), which corresponds in size to the mature processed native STAR protein, was nonfunctional. Confocal microscopy further confirmed that truncation of the minimal N-terminal sequence (^ΔN36-^STAR) abolished mitochondrial localization, mirroring the cytoplasmic distribution observed with the complete STAR*^MTS^* deletion (^ΔN62-^STAR) (Fig 1F). This result strongly suggests that mitochondrial targeting is essential for STAR’s biological activity.

These observations demonstrate that the STAR*^MTS^* is indispensable for mitochondrial cholesterol import and steroidogenesis, challenging previous models that proposed STAR functions independently of mitochondrial entry, and is fully functional without the MTS (*5*, *7*). The discrepancy between our physiological model and the F2-based COS-1 cell system highlights potential artifacts in prior interpretations of STAR’s mechanism of action (*2*, *3*, *5*, *7*, *8*, *22*), emphasizing the relevance of models that represent physiological *de novo* steroid biosynthesis.

### There is no steroidogenic activity for STAR outside mitochondria

Previous studies using the F2 system in COS-1 cells have suggested that STAR could activate steroidogenesis when anchored externally to mitochondria, without requiring mitochondrial entry. Specifically, a fusion protein of ^ΔN62-^STAR tethered to the outer mitochondrial membrane (OMM) via TOM20 was reported to achieve maximal and constitutive steroid production exceeding that of native STAR (*5*). To rigorously test this claim, we expressed the same TOM20-^ΔN62-^STAR fusion protein in MA-10*^STKO^* cells. In contrast to earlier reports, we observed no steroidogenic activity with the TOM20-^ΔN62-^STAR construct (Fig 2A). These findings strongly indicate that mitochondrial entry of STAR is essential for its steroidogenic function.

**Fig. 2.**
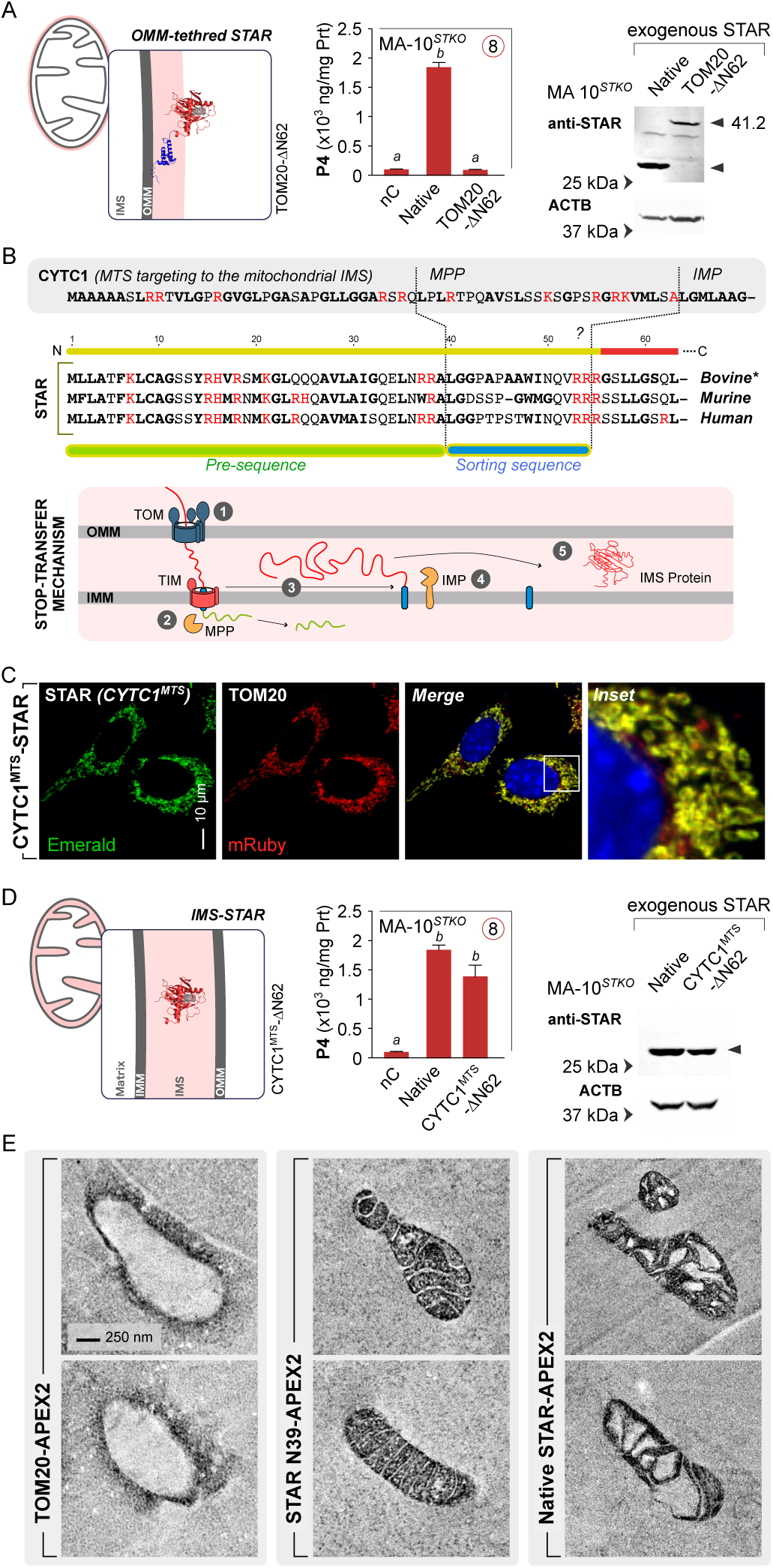
STAR requires import into the mitochondrial intermembrane space (IMS) for function. (**A**) Lentiviral expression of a truncated STAR lacking its mitochondrial targeting sequence (^ΔN62-^STAR) fused to the outer mitochondrial membrane (OMM) anchor TOM20 fails to rescue Bt2cAMP-induced steroidogenesis in STAR-deficient MA-10*^STKO^* cells, in contrast to reconstitution with native STAR (positive control, pC; negative control, nC; different alphabets indicate significant difference p<0.05). Immunoblot confirms expression of the TOM20^-ΔN62-^ STAR fusion protein at the expected molecular weight (∼41.2 kDa). This contrasts with previous claims that this construct yields maximal constitutive steroid production. **(B)** Sequence comparisons between the STAR^MTS^ and the well-characterized bipartite MTS of cytochrome C1 (CYTC1^MTS^) reveal conserved features, including a mitochondrial processing peptidase (MPP) cleavage site and a predicted inner membrane peptidase (IMP) site. These conserved elements support a stop-transfer import mechanism characteristic of proteins targeted to the mitochondrial intermembrane space (IMS). The accompanying schematic illustrates the principle of this stop- transfer mechanism, where translocases at the outer and inner mitochondrial membranes mediate import ①, and MPP-mediated cleavage ②, the sorting signal anchors to the IMM ③, followed by IMP-mediated cleavage ④, and IMS release ⑤. **(C)** Confocal imaging demonstrates mitochondrial colocalization of CYTC1^MTS-ΔN62-^STAR-Emerald with the OMM marker TOM20- mRuby in MA-10 cells, confirming successful targeting to mitochondria using the CYTC1^MTS^. **(D)** Functional rescue of steroidogenesis is achieved in MA-10*^STKO^* cells by CYTC1^MTS-ΔN62-^ STAR, showing equivalence to native STAR in restoring progesterone production, thereby demonstrating that IMS localization fully restores STAR function (different alphabets indicate significant difference p<0.05). Western blot analysis shows identical of mitochondrial processing of native STAR and CYTC1^MTS-ΔN62-^STAR. **(E)** APEX2-based ultrastructural labeling distinguishes STAR sub-mitochondrial localization: TOM20-APEX2 marks the cytoplasmic face of the OMM; ^N36-^STAR-APEX2, containing only the presequence, targets the matrix; native STAR-APEX2 localizes specifically to the IMS. This confirms that the full STAR^MTS^ directs import via a stop-transfer mechanism, releasing STAR into the IMS for its functional role.

### STAR is targeted to and functions within the mitochondrial intermembrane space

Although a previous study on post-translational processing of bovine STAR identified the cleavage of two peptides from the STAR*^MTS^*(*26*), this was deemed inconsequential for its steroidogenic function in COS-1 cells using the F2 approach. We reinterpreted this cleavage pattern as indicative of a bipartite signal sequence, characteristic of Class I mitochondrial intermembrane space (IMS) proteins such as cytochrome C1 (CYTC1) (*27*). Supporting this interpretation, conserved cleavage sites corresponding to a mitochondrial processing peptidase (MPP) and a putative inner membrane peptidase (IMP) were identified in bovine, human, and murine STAR*^MTS^*, closely resembling the well-characterized IMS import system seen in the CYTC1*^MTS^* (Fig 2B). As in other Class I IMS proteins, this configuration comprises an initial matrix-targeting sorting signal followed by a stop-transfer sorting signal that engages translocases at both the OMM and IMM, enabling cleavage and release of the protein into the IMS (*28*).

To functionally test our hypothesis, we engineered a construct in which the mitochondrial targeting sequence of cytochrome C1 (CYTC1*^MTS^*) was fused to the N-terminus of ^ΔN62-^STAR, thereby redirecting ^ΔN62-^STAR to the mitochondrial IMS. Confocal imaging in MA-10 cells confirmed that CYTC1*^MTS^*^-ΔN62-^STAR localized to mitochondria (Fig 2C). When expressed in MA-10*^STKO^* cells, this construct fully restored steroidogenesis, matching the activity of native STAR (Fig 2D). Immunoblot analysis revealed proper post-translational processing of the CYTC1*^MTS^*^-ΔN62-^STAR fusion, with a band pattern indistinguishable from that of native STAR. These findings provide direct functional and biochemical evidence that STAR exerts its activity from within the mitochondrial IMS.

To precisely define the sub-mitochondrial localization of STAR, we employed ultrastructural labeling using STAR-APEX2 fusion constructs expressed in MA-10 cells. Native STAR-APEX2 localized specifically to the mitochondrial IMS (Fig 2E). In contrast, TOM20-APEX2, used as a control, produced membrane-bound staining on the cytoplasmic face of the OMM, while ^N36-^ STAR-APEX2, containing only the first segment of the bipartite targeting sequence, localized to the mitochondrial matrix. These results validate the functional features of the STAR*^MTS^* as a bipartite targeting signal and further corroborate that native STAR is an IMS-resident protein.

### STAR functions as an IMS cholesterol shuttle

A long-standing conceptual challenge in the field has been how STAR, presumed to act at the cytoplasmic face of the OMM, could facilitate high-volume cholesterol transfer with a presumed 1:1 binding stoichiometry for the START domain. This difficulty gave rise to the “molten globule” model, which proposed that STAR remains in a partially unfolded, metastable state outside the mitochondria to perform its function (*16*). However, the biophysical basis of this model lacks direct support from cellular systems. For example, it has been suggested that the acidic pH (∼3.5) required to induce the molten globule transition is a proxy for the mitochondrial membrane potential (ΔѱM), thereby linking STAR activation to mitochondrial bioenergetics. Yet, the ΔѱM is a prerequisite for all mitochondrial protein import (*29*), not a specific determinant for STAR folding/unfolding or function.

The association of STAR with mitochondrial import components such as TOM (*18*) and TIM (*30*) complexes, previously interpreted as part of a specialized cholesterol import complex (*16*), is more plausibly explained by the necessity for STAR to enter the intermembrane space (IMS) via canonical protein import pathways. This reframing places the focus on STAR as a typical IMS protein, rather than a cytoplasmic actor reliant on partner proteins at the outer membrane.

Independent lines of evidence have hinted at an alternative cholesterol shuttle mechanism. Early reconstitution studies using liposomal systems demonstrated that STAR can transfer cholesterol between membranes in a manner similar to sterol carrier protein 2 (SCP2), supporting its potential as a lipid shuttle (*31*). Structural comparisons of START domain-containing proteins further suggested a conformation well-suited for cholesterol binding and transport in a folded state (*32*). Although these models were historically overlooked in the steroidogenesis field (*33*), they underscore that the START domain of STAR is structurally optimized for cholesterol trafficking without requiring unfolding into a molten globule.

To investigate whether STAR possesses structural features consistent with a shuttle mechanism, we used PPM 3.0 (*34*), a membrane orientation prediction tool, to analyze human STAR’s spatial positioning of proteins relative to lipid bilayers. This analysis identified two flexible surface loops – Loop 7 and Loop 11 – as containing conserved hydrophobic residues capable of embedding into membranes. Specifically, L177 in Loop 7 and W249 and P251 in Loop 11 were predicted to anchor within the membrane bilayer (Fig. 3A). These results are consistent with a recent structural study indicating that Loop 7 flexibility underlies lipid transfer function of START domain proteins (*35*).

**Fig. 3.**
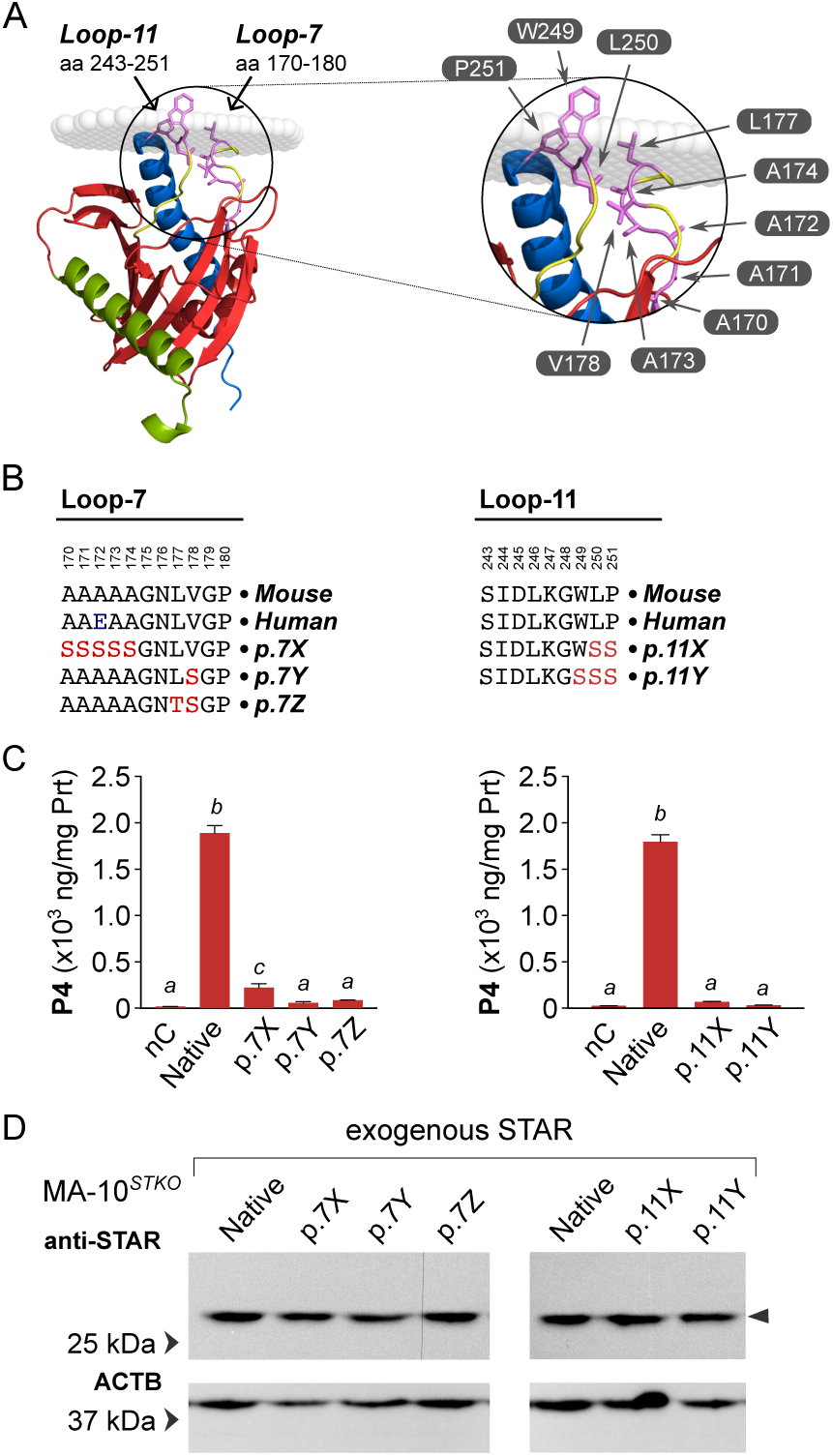
Membrane association of STAR is essential for its cholesterol transfer function. Membrane orientation modeling of human STAR identified key conserved hydrophobic residues predicted to mediate membrane docking of STAR. In Loop 7, L177 is embedded in the membrane, flanked by V178 and residues A170–A174. In Loop 11, W249 and P251 are membrane-embedded, with L250 occupying an intermediate position. These two flexible surface loops are structurally positioned to enable transient membrane interactions necessary for lipid transfer. **(B)** Site-directed mutagenesis was used to disrupt these hydrophobic interactions. Loop 7 mutants included: A170S-A171S-A172S-A173S-A174S (p.7X), L177T- V178S (p.7Y), and V178S alone (p.7Z); amino acid numbers are based on the murine sequence. Loop 11 mutants included: L250S-P251S (p.11X) and W249S-L250S-P251S (p.11Y). These constructs were used to assess STAR function in MA-10*^STKO^* cells. **(C)** None of the Loop 7 or Loop 11 mutants were able to restore progesterone production in MA-10*^STKO^* cells, in contrast to native STAR, demonstrating the functional requirement for these hydrophobic residues in membrane-associated cholesterol shuttling (negative control, nC; different alphabets indicate significant difference p<0.05). **(D)** Western blot analysis confirmed comparable expression levels of all mutant and native STAR proteins, indicating that the loss of function was not due to differences in protein stability or expression efficiency.

To test the functional importance of these hydrophobic residues, we performed site-directed mutagenesis to substitute them with hydrophilic amino acids. In MA-10*^STKO^* cells, expression of Loop 7 mutants [p.7X(A170S-A174S), p.7Y(V178S), and p.7Z(L177T-V178S)] and Loop 11 mutants [p.11X(L250S-P251S) and p.11Y(W249S-L250S-P251S)] failed to rescue steroidogenic function (Fig. 3C-D). All mutants were expressed at comparable levels to wild-type STAR, as confirmed by immunoblotting, indicating that their loss of function was not due to instability or degradation.

These results collectively demonstrate that STAR requires direct, flexible interaction with membranes for function, mediated through hydrophobic residues in Loops 7 and 11. The data support a model in which folded STAR, localized to the mitochondrial IMS, functions as a cholesterol shuttle by transiently docking onto membranes via these loops to extract and deliver cholesterol to the inner mitochondrial membrane. This mode of action stands in contrast to previous models involving unfolded intermediates and places STAR squarely within the established framework of IMS protein import and lipid transfer.

### STAR degradation is via mitophagy

The molten globule model posited that STAR completes its function *en route* to the mitochondrial matrix (*5*), thereby implicating Lon proteases within the matrix as being responsible for its degradation (*36*, *37*). However, this narrowly focused interpretation was not based on direct evidence and alternative possibilities were largely overlooked. Recognizing that STAR is folded and functions within the mitochondrial intermembrane space (IMS), we reasoned that its entry into the matrix, where Lon proteases reside, is highly improbable. Instead, we hypothesized that STAR is degraded through cytosolic quality control mechanisms.

Supporting this idea, earlier studies had observed extensive autophagy in steroidogenic cells, speculated to be linked to the regulation of steroid levels (*38*), prompting us to investigate mitophagy as a potential mechanism for rapid STAR degradation.

To test this hypothesis, we inhibited mitophagy by blocking mitochondrial fission using MDIVI-1, a DRP1 inhibitor (*39*). In MA-10 cells, MDIVI-1 treatment effectively prevented STAR degradation following steroidogenic stimulation (Fig. 4A). Notably, STAR protein levels remained elevated even after removal of the stimulatory signal, and steroid production remained persistently high. This indicated that preventing mitochondrial fission arrests STAR at its IMS site of action and halts its degradation. Similarly, inhibition of lysosomal acidification using the proton pump inhibitor lansoprazole (*40*) also blocked STAR degradation (Fig. 4A), further implicating lysosome-dependent mitophagy as the degradation pathway.

**Fig. 4.**
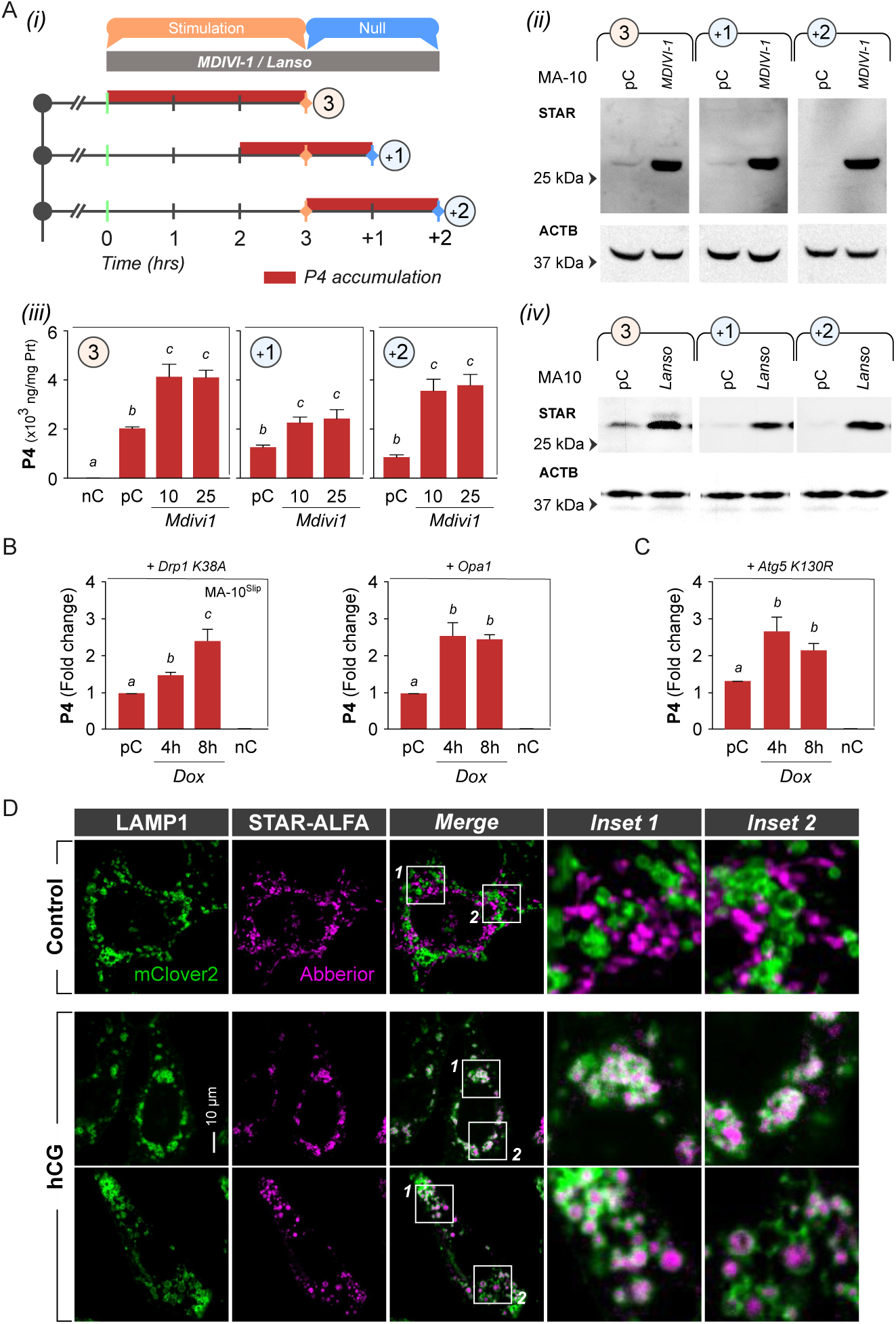
STAR is degraded via mitophagy. (**A**) Pharmacological inhibition of mitochondrial fission using MDIVI-1, a DRP1 inhibitor, prevents STAR degradation. (i) Experimental design showing period of Bt2cAMP stimulation and withdrawal (null) timepoints in MA-10 cells. (ii) Western blots show elevated STAR protein levels at 3 hours of stimulation, and persistence of STAR protein after 1 hour and 2 hours of null, compared to untreated controls (pC). (iii) Progesterone (P4) levels remain elevated with MDIVI-1 treatment, particularly evident after medium replacement during null, indicating that fission is required for STAR degradation and termination of steroidogenesis (different alphabets indicate significant difference p<0.05). (iv) Blocking lysosomal acidification with lansoprazole (Lanso) similarly prevents STAR degradation and sustains steroidogenesis, confirming a lysosome-dependent degradation mechanism. **(B)** Genetic inhibition of mitochondrial fission through doxycycline (Dox)-inducible expression of dominant-negative DRP1^K38A^ or mitochondrial fusion protein OPA1 also leads to increased P4 production following stimulation, validating the role of mitochondrial dynamics in regulating STAR turnover (negative control, nC; positive control, pC; different alphabets indicate significant difference p<0.05). **(C)** Inhibition of autophagosome formation by Dox- inducible expression of dominant-negative ATG5^K130R^ likewise increases stimulated P4 production (different alphabets indicate significant difference p<0.05). This supports a two-step degradation mechanism for STAR involving mitochondrial fission followed by autophagy/mitophagy. **(D)** Confocal imaging of STAR-ALFA (a functional C-terminally ALFA- tagged STAR fusion protein; also see Fig S3) and LAMP1-mClover2 (lysosomal marker) reveals no colocalization in unstimulated MA-10 cells. Upon stimulation, STAR-ALFA extensively co- localizes with LAMP1-labeled lysosomes, providing direct visual evidence that STAR is targeted for lysosomal degradation via mitophagy.

We corroborated these findings using two independent genetic strategies to suppress mitochondrial fission: expression of a dominant-negative DRP1 mutant (DRP1^K38A^) (*41*) and overexpression of the mitochondrial fusion protein OPA1 (*42*). Both approaches resulted in elevated STAR activity, reflected by sustained steroidogenesis in MA-10 cells (Fig. 4B). Additionally, blocking autophagosome formation through overexpression of the dominant-negative autophagy mutant ATG5^K130R^ similarly enhanced steroid production (Fig. 4C), consistent with a two-step process involving mitochondrial fission followed by autophagy-mediated degradation of STAR.

To visualize mitophagy directly, we used LH-responsive MA-10*^Slip5^*Leydig cells (*43*) expressing TOM20-mRuby to label mitochondria and LAMP1-mClover2 to label lysosomes. Upon trophic stimulation with hCG, we observed pronounced co-localization of mitochondria and lysosomes (Fig. S2), a pattern absent in unstimulated controls and consistent with the clearance of mitochondrial regions via mitophagy. To assess STAR localization during this process, we expressed a C-terminal ALFA-tagged STAR (STAR-ALFA), which fully rescued steroidogenesis in MA-10*^STKO^* cells (Fig. S3). In unstimulated cells, STAR-ALFA did not co- localize with LAMP1-labeled lysosomes. However, upon stimulation, we observed robust co- localization of STAR-ALFA with LAMP1-mClover2 (Fig. 4D), directly indicating that STAR is targeted for degradation through mitophagy.

### Interpretations for human STAR mutations in lipoid CAH

To date, numerous mutations in the human STAR gene have been implicated in lipoid congenital adrenal hyperplasia (CAH), with varying severity of clinical phenotypes (Table S1 and S2; Fig 5A). However, the mechanistic basis for loss-of-function in many of these mutations has remained unclear. In particular, mutations affecting the STAR*^MTS^* have largely been reported only in the context of frameshift mutations or premature stop codons, leading to the impression that isolated, in-frame mutations within the STAR*^MTS^* do not contribute to pathological phenotypes (*7*). This notion warrants re-evaluation in light of both our findings and a recent clinical report describing a lipoid CAH patient with an in-frame deletion of amino acids 22–59 within the STAR*^MTS^* (*44*). The functional deficit in this case is readily explained by impaired mitochondrial targeting, preventing STAR from reaching its IMS site of action.

**Fig. 5.**
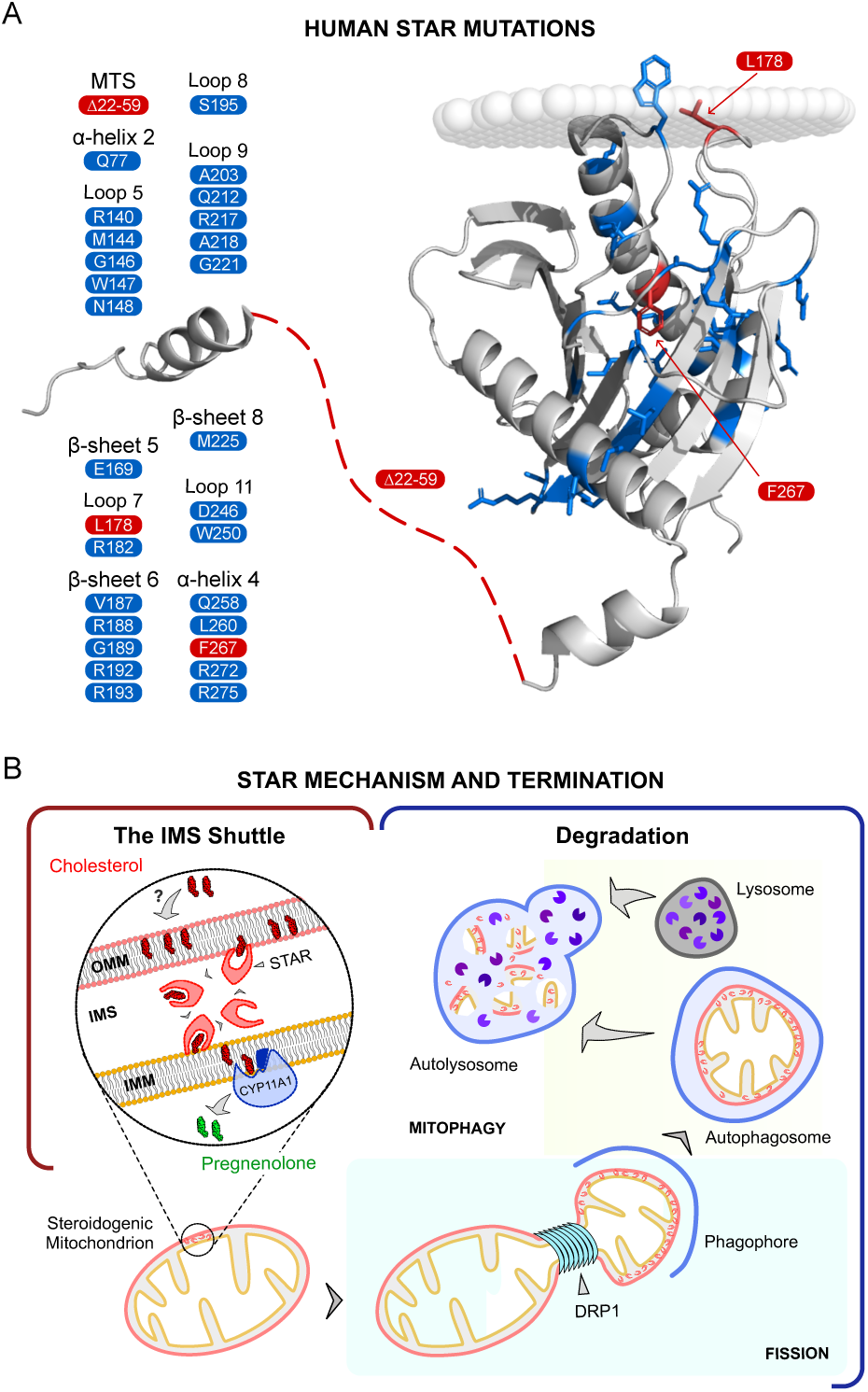
Integration of STAR’s mechanism of action with clinical mutation mapping. (**A**) Structure of human STAR showing disease-associated mutations reported in patients with various severities of lipoid congenital adrenal hyperplasia (CAH). Mutations span the MTS, membrane-interacting loops (Loop 7, 9, 11), and the C-terminal α-helix. These regions correspond to the functional elements identified in this study (shown in Red; See Tables S1 and S2 for mutation details). **(B)** Schematic summarizing STAR function and turnover. STAR is imported into the mitochondrial IMS via a stop-transfer mechanism, where it acts as a cholesterol shuttle. Cholesterol is delivered to the IMM, where CYP11A1, localized to the matrix-facing IMM surface, converts it into pregnenolone, the first steroid in *de novo* steroidogenesis. STAR-enriched mitochondrial subdomains are subsequently cleared via mitophagy, involving mitochondrial fission, autophagosome formation, and lysosomal fusion into an autolysosome that mediates STAR degradation.

Moreover, mutations outside the canonical cholesterol-binding pocket of the START domain have remained poorly understood. For example, a previously reported L178Q mutation, associated with lipoid CAH (*45*) (Fig 5A), maps to Loop 7, a region we identified as critical for membrane interaction (Fig 3). Similarly, the C-terminal mutation F267S (*46*), also linked to lipoid CAH, affects a hydrophobic residue in the final α-helix of STAR (Fig 5A, Fig S4), which we showed to be important for proper START domain function.

By establishing the essential roles of the STAR*^MTS^*, membrane-interacting loops and C-terminal hydrophobic residues outside the START domain, our findings elucidate the principle by which cholesterol is transferred from the outer to inner mitochondrial membrane. This provides mechanistic clarity for a broader spectrum of pathogenic STAR mutations. These insights not only expand the framework for interpreting patient genotypes but also reinforce the structural basis underlying mitochondrial cholesterol import in steroidogenesis.

## DISCUSSION

From the time Orme-Johnson and colleagues first described the rapid induction of a protein in response to steroidogenic stimulation (*47*) – later characterized as STAR (*1*) – identifying its precise mechanism of action became crucial for both understanding steroidogenesis and addressing the pathology of lipoid CAH in patients (*22*). Despite extensive investigations, the field has faced significant challenges due to competing models, particularly concerning the purported involvement of TSPO (*48*), and conflicting experimental approaches utilizing non-physiological systems. Our present work provides definitive clarity on STAR’s cellular localization, function, and degradation, resolving key aspects of these controversies.

Historically, models suggesting that STAR operated externally to mitochondria, potentially through interactions with outer mitochondrial membrane proteins like TSPO, gained significant traction (*8–10*). However, our previous findings, using rigorous genetic approaches, clearly negated the necessity of TSPO for mitochondrial cholesterol import (*11–13*), thereby necessitating a reevaluation of STAR’s mechanism without preconceived associations. Our current data, derived from a physiologically relevant steroidogenic MA-10 Leydig cell system with endogenous steroidogenic capacity, demonstrates unequivocally that STAR is an IMS-localized cholesterol shuttle rather than a cytoplasmic tether. Our results thus challenge longstanding models proposing cytoplasmic or outer mitochondrial localization and molten-globule transitions (*16*).

The significance of the mitochondrial targeting sequence (MTS) of STAR had previously been dismissed based on observations from COS-1 cell-based synthetic systems (*6*, *7*). In contrast, our findings emphasize the critical importance of the STAR MTS in proper mitochondrial localization and function. The exclusive functional rescue achieved by targeting STAR specifically to the intermembrane space (IMS) using the CYTC1^MTS^ fusion unequivocally establishes the IMS as STAR’s physiological site of action and provides a direct mechanistic explanation for a human disease-causing mutation within the MTS region (*44*). Moreover, structure-function analyses, informed by modeling and mutagenesis, identified specific hydrophobic loops essential for cholesterol binding and transfer, providing molecular insights consistent with pathological human mutations identified in lipoid CAH patients (*45*, *46*).

With the new understanding from this study that STAR localizes specifically to the mitochondrial IMS and functions as a cholesterol shuttle, previous suggestions of macromolecular complexes containing STAR or restricted protein movements and folding (*49*), appear unlikely. Furthermore, earlier interpretations involving chaperones such as GRP78 (*17*) or mitochondrial import machinery components like TOM40 (*18*) and TIM50 (*30*) must now be reconsidered. Rather than participating directly in mitochondrial cholesterol import, these factors likely facilitate the import of STAR into the IMS, indirectly influencing steroidogenesis.

Although the potential functional importance of the STAR^MTS^ in steroidogenesis was previously suggested through an *in vivo* transgenic mouse model expressing ^ΔN47-^STAR in a STAR knockout background (*50*), conflicting outcomes from *in vitro* studies employing the synthetic F2 system led to uncertainties regarding its role. Our results now clarify this apparent discrepancy, providing a robust mechanistic explanation for the disrupted steroidogenic function observed in the earlier *in vivo* experiments.

Our identification of mitophagy as a rapid degradation mechanism for STAR provides a novel insight into steroidogenic regulation. Previously proposed models focused largely on mitochondrial matrix proteases, particularly Lon protease (*36*, *37*). However, our comprehensive pharmacological and genetic data—including the inhibition of mitochondrial fission (via DRP1 inhibition) and autophagosome formation (via ATG5 dominant-negative expression)—strongly support mitophagy as a primary pathway for STAR degradation. This discovery not only resolves prior ambiguities regarding the temporal regulation of steroidogenesis but also implicates a broader relevance of mitophagy in the regulation of IMS proteins. Furthermore, our findings provide an accurate biological context for earlier ultrastructural observations describing robust autophagic activity accompanying steroidogenesis (*38*).

While our study significantly advances the understanding of STAR’s precise localization, function, and regulated degradation, we recognize certain considerations for future investigation. One notable area is the mechanism by which STAR-containing mitochondrial regions are selectively targeted for elimination. Although we have shown that mitophagy is responsible for STAR degradation, the specificity of this process, how only certain mitochondrial domains harboring STAR are marked for removal while others might be spared, remains unclear. This raises intriguing questions about mitochondrial compartmentalization and quality control, and identifying the molecular signals or conditions that drive this selectivity will be an important direction for future research. Additionally, while the functional consequences of hydrophobic loop mutations in STAR were rigorously tested through reconstitution in a physiological knockout system, our conclusions were drawn primarily from cellular rescue assays. Direct biochemical measurements of cholesterol binding and membrane association for these mutants were not performed. Although the strong concordance between our experimental findings and clinical phenotypes observed in patients with analogous human STAR mutations provides independent support, future structural and biochemical studies would help validate and refine the mechanistic details of how STAR engages with mitochondrial membranes. Such work could also reveal additional nuances of START domain function and dynamics that underlie cholesterol transfer activity.

In summary, our findings provide a cohesive and mechanistically precise reinterpretation of STAR-mediated mitochondrial cholesterol import, overcoming historical controversies and misinterpretations in the field. By establishing STAR as a mitochondrial IMS cholesterol shuttle subject to rapid degradation via mitophagy, we have identified a novel milestone in understanding protein degradation within the mitochondrial intermembrane space, particularly in an extremely acute and functionally regulated biological system. These insights fundamentally reshape our understanding of steroidogenic regulation and offer direct translational relevance for diagnosing and interpreting clinical outcomes associated with STAR mutations. Moving forward, our study sets a robust foundation for further exploration into mitochondrial cholesterol metabolism, with significant implications across the fields of endocrinology, metabolism, and mitochondrial biology.

## MATERIALS AND METHODS

### Cells and steroidogenesis

The MA-10 Leydig cell line (*51*), used in the original characterization of STAR protein (*1*), served as the steroidogenic model system. Cells were cultured at 37°C in a humidified incubator with 5% CO2, in DMEM high glucose medium supplemented with 1 mM sodium pyruvate, 10% fetal bovine serum, 1% non-essential amino acids, and 1% penicillin-streptomycin. For induction of steroidogenesis, cells were treated with either 0.5 mM dibutyryl cAMP (Bt2cAMP) or 1.5 IU/ml human chorionic gonadotropin (hCG) for 3 hours. Progesterone levels in culture supernatants were quantified using radioimmunoassay (RIA), as previously described (*12*). All experiments were conducted with three biological replicates and repeated independently at least three times.

### STARD1 knockout MA-10 cells

CRISPR/Cas9 gene editing was used to target exon 2 of the murine STAR locus (Gene ID: 20845), creating null alleles in STAR knockout MA-10 cell (MA-10*^STKO^*) clones. In brief, the targeting guide RNA (5′-GGATGGGTCAAGTTCGACGT-3′) was cloned into the pSpCas9(BB) vector (*52*) and transfected into MA-10 cells. Single-cell clones were established in 96-well plates at limiting dilution and expanded. Clones were screened for STAR protein by immunoblotting. Functional knockout was confirmed by absence of progesterone synthesis after Bt2cAMP stimulation, with full rescue of steroidogenesis in the presence of 22- hydroxycholesterol, indicating a specific defect in mitochondrial cholesterol import/STAR.

### Immunoblotting

Western blot detection of endogenous and transgenic protein expression in MA-10 cell lysates has been previously described (13, 38). In brief, cells were lysed in Laemmli sample buffer supplemented with protease inhibitors and disrupted by sonication. Proteins were denatured at 95°C for 5 minutes, separated by SDS-PAGE, and transferred to PVDF membranes. STAR protein was detected using a polyclonal rabbit anti-STAR antibody (*43*) or FluoTag®-X2 anti-ALFA (NanoTag Biotechnologies). Chemiluminescent detection was performed with Poly- HRP-conjugated secondary antibodies. An IRDye800-conjugated β-actin antibody (LI-COR Biosciences) served as a loading control. Imaging was performed using a C600 imager (Azure Biosystems).

### Hormone assays

Progesterone levels were measured using a previously validated radioimmunoassay (RIA) (*12*). For each condition, medium was collected following the indicated treatment duration and clarified by centrifugation. Samples were stored at -20°C until assayed. RIA was performed using tritiated progesterone and a polyclonal antibody with high specificity for progesterone (*53*). Standard curves were generated for each assay, and sample concentrations were calculated by interpolation. Hormone levels were normalized to total protein content measured by BCA assay.

### STAR structure

Structural features of the murine STAR protein (NCBI mRNA: NM_011485.5) were analyzed using a combination of experimental and computational approaches. The human STAR crystal structure (PDB: 3P0L) was used as the primary reference for conserved elements of the START domain, including the hydrophobic ligand-binding pocket and surrounding α-helices and β-sheets. Murine STAR structure was modeled using AlphaFold (AF-Q60920-F1), and structural alignments were performed in PyMOL to compare murine and human STAR and highlight conserved residues.

### Vector and transgene designs

All vectors and transgenes designed for this project are available Table S3. Native murine STAR (NCBI mRNA: NM_011485.5) was cloned using RT-PCR in a pMX retroviral vector (pMX-hOCT4; Addgene plasmid 17217) to generate pMX-STAR. This sequence confirmed plasmid was used as a template to generate truncated, fusion and mutated STAR proteins. The N-terminal 36, 42, 50, 54 and 62 amino acids were truncated by designing forward primers starting at nucleotides 106, 127, 151, and 163 from the 5’ end of the native STAR (designated ΔN36, ΔN42, ΔN50, ΔN54 and ΔN62-STAR). To ensure successful ribosomal assembly, the Kozak sequence including start codon, ATG were added to the forward primers for the N-terminal truncations. The C-terminal deletion of 10 and 24 amino acids were performed by reverse primers designed at nucleotides 33 and 75 from 3’ end respectively (designated as ΔC10 and ΔC24). To ensure successful termination, a stop codon was incorporated in the reverse primers for the C-terminal deletions.

Fusion of fluorescent proteins were also constructed in the pMX retroviral vector, for which a C-terminal Emerald (from Emerald-N1; Addgene plasmid 54588) was inserted into pMX-ΔN36-STAR and pMX-ΔN62-STAR. Fluorescent proteins targeting the OMM was constructed by fusion of TOM20 (amplified from murine testis cDNA) and fluorescent protein mRuby (from mRuby-N1; Addgene plasmid 54581) in the pMX retroviral vector. TOM20 was also inserted at the N-terminal of ΔN62-STAR (in the pMX-ΔN62-STAR plasmid) to yield pMX-TOM20-ΔN62-STAR. The CYTC1 MTS (N-terminal 84 amino acids, amplified from murine testis cDNA), was also inserted at the N-terminal of ΔN62-STAR (in the pMX-ΔN62 plasmid) to yield pMX-CYTC1-MTS-ΔN62-STAR. For localization, the CYTC1-MTS was also fused with a C-terminal Emerald (pMX-CYTC1-MTS-Emerald).

Site-directed mutagenesis was performed in pMX-STAR by designing long primers flanking the target nucleotides in STAR followed by extension across the vector and template digestion using DpnI. The Loop 7 amino acids, A170, A171, A172, A173 and A174 in combination were mutated to S170, S171, S172, S173 and S174 (designated as p.7X); L177 and V178 in combination were mutated to S177 and S178 (designated as p.7Z); V178 was mutated to S178 (designated as p.7Y). The Loop 11 amino acids, W249, L250 and P251 in different combinations were mutated to S249, S250 and S251 (designated as p.11X and p.11Y). In the C-terminal α- helix of the START domain, amino acids I264, F266, A267 and L270 in combination were mutated to T264, W266, S267 and R270 (designated as p.⍺4).

Fusions of STAR with an engineered ascorbate peroxidase (APEX2) for ultrastructural localization was cloned from mito-V5-APEX2 (72480; Addgene) downstream of native and N36-STAR in the pMX retroviral vector (pMX-STAR-APEX2, pMX-N36-APEX2). For OMM labeling, a pMX-TOM20-APEX2 was also constructed.

For fluorescent protein labeling of lysosomes, LAMP1-mClover2 (from Clover-Lysosomes- 20; Addgene plasmid 56528) was cloned into the pMX retroviral vector (pMX-LAMP1- mClover2). For STAR immunolocalization, a synthetic ALFA-tag was added to the C-terminal of STAR and cloned in a doxycycline-inducible lentiviral vector, pLenti-TRE-rtTA (*54*), to yield a pLenti-TRE-STAR-ALFA-EF1⍺-rtTA vector. For preventing mitochondrial fission OPA1 (from pMSCV-OPA1; Addgene plasmid 26407), and DRP1^K38A^ (from pBABE-puro-hDrp1- K38A; Addgene plasmid 37243), were cloned into pRRL-TRE-rtTA (pRRL-TRE-OPA1-PGK-rtTA and pRRL-TRE-DRP1^K38A^-PGK-rtTA). Similarly for dominant negative inhibition of autophagy ATG5^K130R^ (from pmCherry-ATG5-K130R; Addgene plasmid 13096), was cloned into pRRL-TRE-rtTA (pRRL-TRE-ATG5^K130R^-PGK-rtTA).

### Viral packaging and transductions

Expression of the different STAR mutations, fusions and fluorescent proteins were done using retroviral or lentiviral systems. Packaging and infections for both lentiviruses and retroviruses has been described in previous work (*55*, *56*). In brief, retroviral packaging was performed in GP2 cells with co-transfection of a helper plasmid pCMV-VSV-G (Addgene plasmid 8454). Lentiviral packaging was performed in 293T cells with co-transfection of vectors psPAX2 (Addgene plasmid 12260) and pMD2.G (Addgene plasmid 12259). Medium collected from packaging cells were cleared by centrifugation and filtration. Infections were carried out in culture medium with added polybrene (10 µg/ml).

### Lysosomal degradation

To evaluate STAR degradation, MA-10 cells were treated with either a DRP1 inhibitor MDIVI-1 (25 µM), to block mitochondrial fission, or the proton pump inhibitor lansoprazole (200 µM) to prevent lysosomal acidification and proteolysis. As per the treatment timelines as specified (Fig 4A), cells were stimulated with Bt2cAMP, followed by withdrawal of stimulation to monitor degradation kinetics. In parallel, genetic approaches expressing a dominant negative form of DRP1 (DRP1^K38A^) or OPA1 were used to inhibit mitochondrial fission. Another dominant negative autophagy mutant ATG5^K130R^ was used to block autophagosome formation. Doxycycline (Dox, 2 µg/ml) was used to induce transgenic protein expression from lentiviral integrations for 4 or 8 hours. In these studies, STAR protein expression was evaluated using Western blots, and progesterone accumulation in the medium was measured using RIA.

### Fluorescent localization

Sample preparations for localization of fluorescent proteins and immunolocalization was performed using methods previously described (*54*), using a confocal microscope (Leica TCS SP5). For fluorescent protein imaging (Emerald, mRuby and mClover2), MA-10 cells were fixed in 4% formaldehyde, rinsed and mounted in antifade mountant (ProLong Gold, Thermo Fisher Scientific). For ALFA-tag immunocytochemistry, fixed cells were permeabilized using 1% Tx- 100 for 1 minute, rinsed and incubated with FluoTag®-X2 anti-ALFA (fluorophore: Abberior STAR 635p), and rinsed prior to mounting.

### Ultrastructural studies

MA-10 cells with transgenic expression of ^N36-^STAR-APEX2, STAR-APEX2 and TOM20- APEX2 were fixed for 60 minutes in 2.5% glutaraldehyde in sodium cacodylate buffer (50 mM, pH 7.4). Fixative was neutralized with 20 mM glycine in cacodylate buffer and rinsed. Samples were then incubated with substrate 3,3’diaminobenzidine (1.4 mM), with 0.03% H2O2 for 20 min. The APEX2 reaction was stopped by washing again in sodium cacodylate buffer, followed by post-fixation overnight at 4°C. Samples were then processed for transmission electron microscopy by dehydration, embedding and preparing 70 nm ultrathin sections (Leica Ultracut UCT). For observation of mitochondrial structure in MA-10 and MA-10*^STKO^* cells, cells were additionally postfixed in osmium tetroxide (1%) for 60 minutes and stained with uranyl acetate (2%) for 60 minutes at 4°C, prior to embedding and sectioning. Mitochondria within cells were imaged using a BioTwin transmission electron microscope (FEI Tecnai 12).

### Statistics

All quantitative experiments were performed using at least three independent biological replicates. Data are presented as mean ± standard error of the mean (SEM). Statistical analyses were conducted using GraphPad Prism v9 (GraphPad Software). For pairwise comparisons, Student’s t-tests were used (including for binary outcomes). For multiple group comparisons, one-way ANOVA followed by Tukey’s post hoc test was applied. A p-value of <0.05 was considered statistically significant. Statistical outcomes are provided in the figure legends. No data points were excluded from the analyses.

## Acknowledgments

The authors extend their heartfelt gratitude to Dr. Douglas Stocco (Texas Tech University Health Sciences Center) for his invaluable feedback and mentorship, and for his unwavering encouragement to pursue scientific truth with rigor and integrity. We also thank John Grazul and Mariena Ramos for their expert assistance with transmission electron microscopy.

## Author contributions

Conceptualization: PPK, AHZ, VS Methodology: PPK, AHZ, VS

Investigation: PPK, AHZ, MCK, ACRF Visualization: PPK, VS

Funding acquisition: VS Project administration: VS Supervision: VS

Writing – original draft: VS, PPK, AHZ Writing – review & editing: VS

## Data and materials availability

All data are available in the main text or the supplementary materials. Expression vectors for the different constructs are available upon request or via addgene.org.

## List of Supplementary Materials

**Fig. S1.**
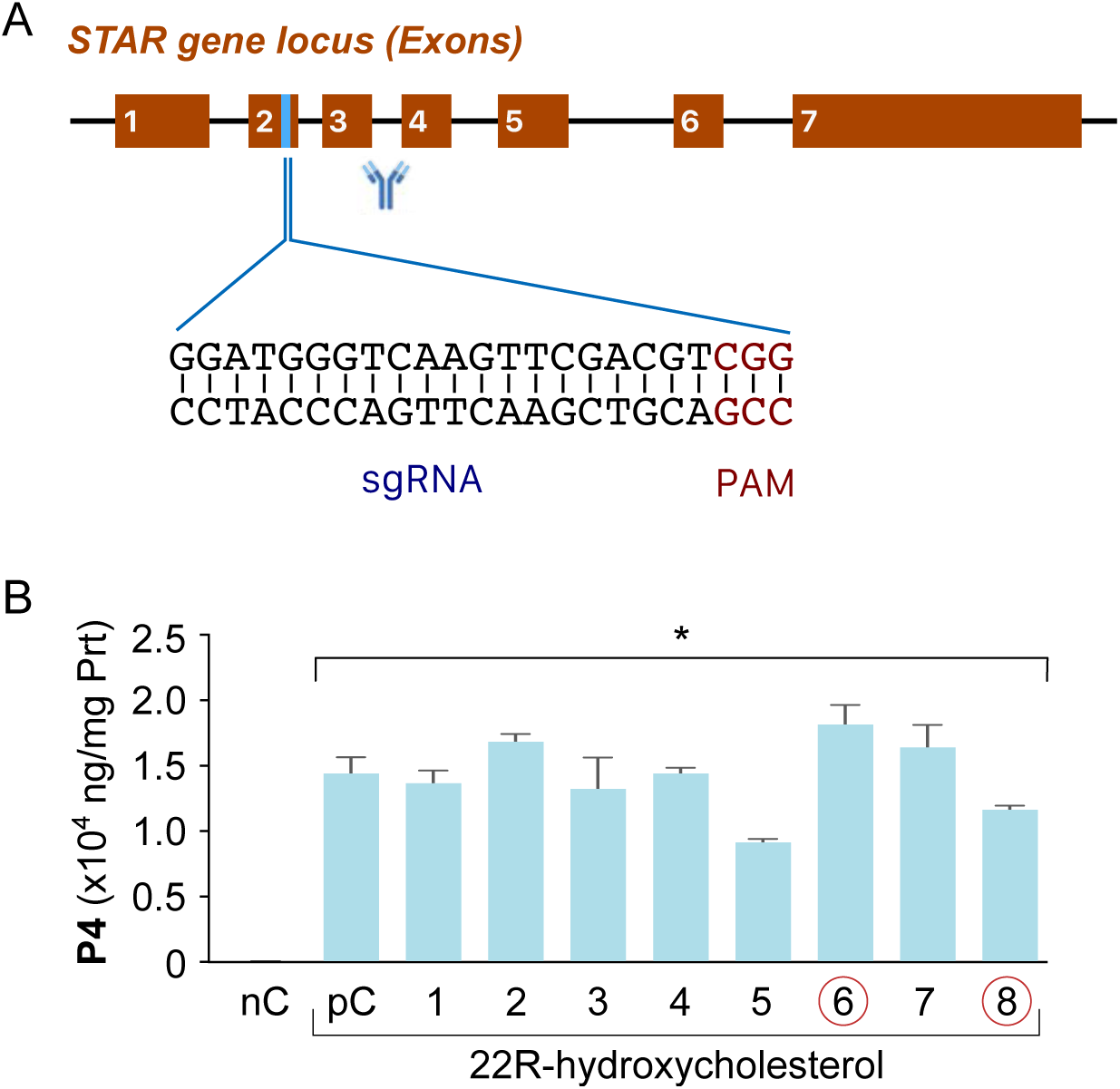
Generation and validation of MA-10 STAR knockout cells. (**A**) Schematic representation of the CRISPR/Cas9 strategy used to delete endogenous STAR. The guide RNA (gRNA) targets exon 2 of the murine STAR genomic locus. The region of the protein detected by the STAR antibody used in immunoblot screening is also indicated. **(B)** Functional validation of STAR knockout clones (MA-10^STKO^). Cells stimulated with 22R-hydroxycholesterol—a water-soluble cholesterol analog that bypasses the need for STAR-mediated mitochondrial import—show robust progesterone (P4) production, confirming intact/native downstream steroidogenic capacity (negative control, nC, positive control, pC; asterisk indicates significant difference from nC, p<0.05). In contrast, these clones fail to produce P4 in response to Bt_2_cAMP stimulation alone (see Fig. 1B), establishing this as a high-fidelity loss-of-function model to test mitochondrial cholesterol import and rescue by STAR constructs.

**Fig. S2.**
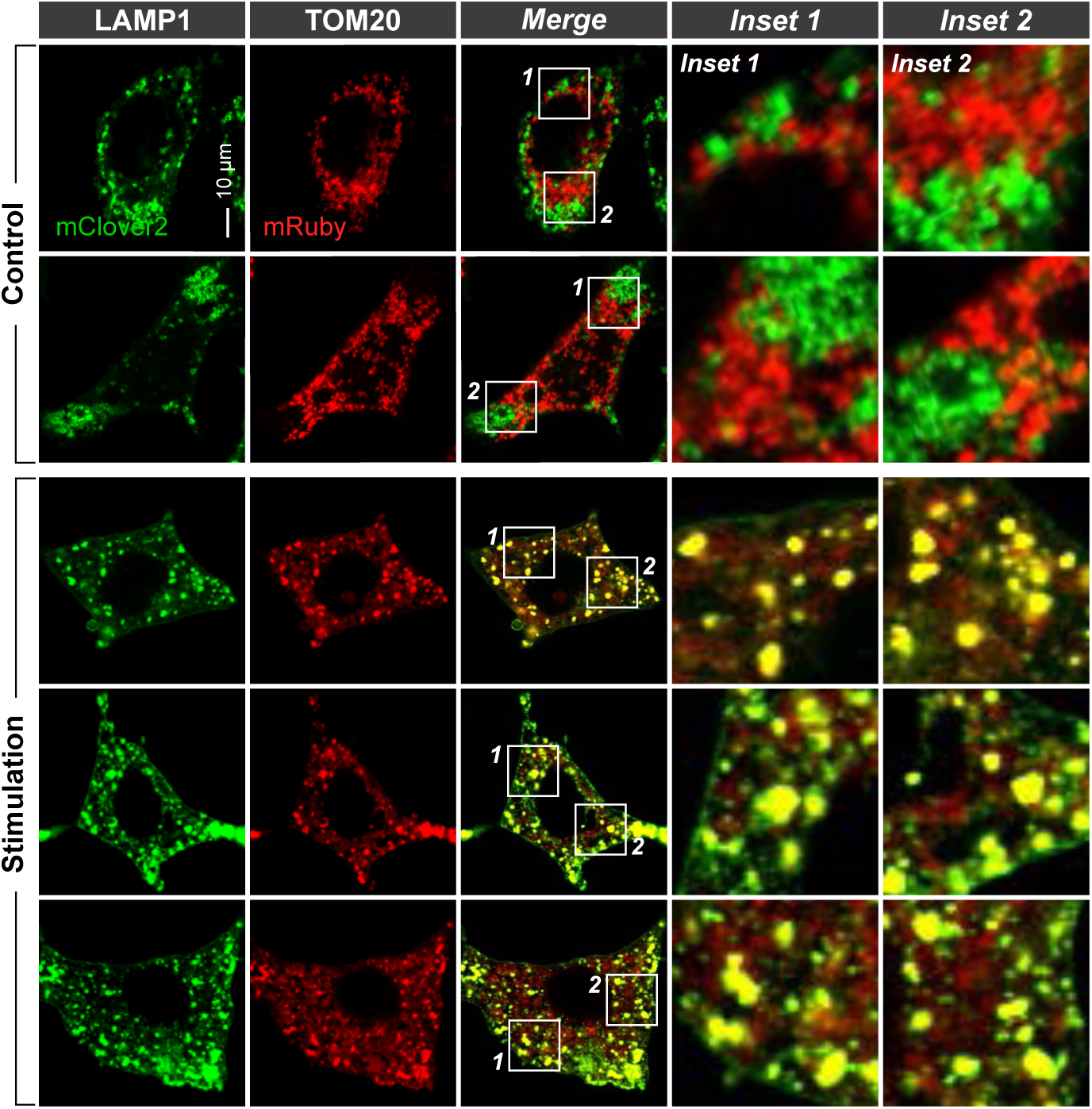
Lysosomal degradation of mitochondria is triggered upon steroidogenic stimulation. Double-labeled MA-10*^Slip5^* cells expressing TOM20-mRuby (mitochondria) and LAMP1- mClover2 (lysosomes) were generated via lentiviral transduction. Confocal imaging of cells under resting conditions (Control) shows no colocalization between mitochondria and lysosomes. In contrast, after stimulation, extensive colocalization of TOM20 and LAMP1 signals is observed, indicating active mitophagy. These findings suggest that mitochondrial subdomains are targeted for lysosomal degradation in response to steroidogenic stimulation, consistent with a mechanism for terminating steroid biosynthesis via mitophagy.

**Fig. S3.**
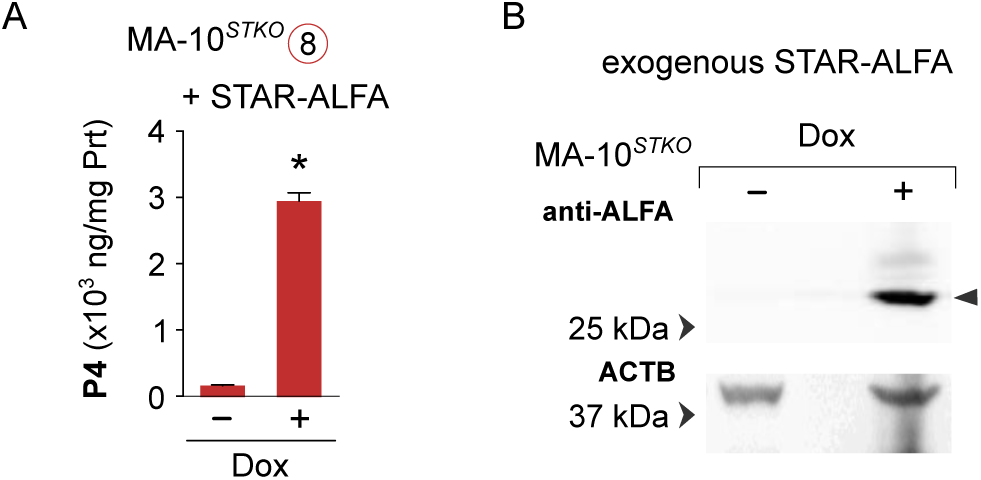
STAR retains full steroidogenic function when fused with a C-terminal ALFA-tag. **(A)** MA10*^STKO^*cells stably expressing a doxycycline (Dox)-inducible STAR-ALFA fusion protein fail to produce progesterone (P4) with stimulation in the absence of Dox induction (–Dox). Upon Dox treatment for 4 hours, STAR-ALFA expression is induced and fully restores steroidogenesis (+Dox) to levels comparable to native STAR, confirming functional integrity of the tagged construct (asterisk indicates significant difference p<0.05). **(B)** Representative Western blot showing robust expression of STAR-ALFA following 4-hour Dox induction, detected using an anti-ALFA nanobody. The loading control (ACTB) confirms equal protein input.

**Fig. S4.**
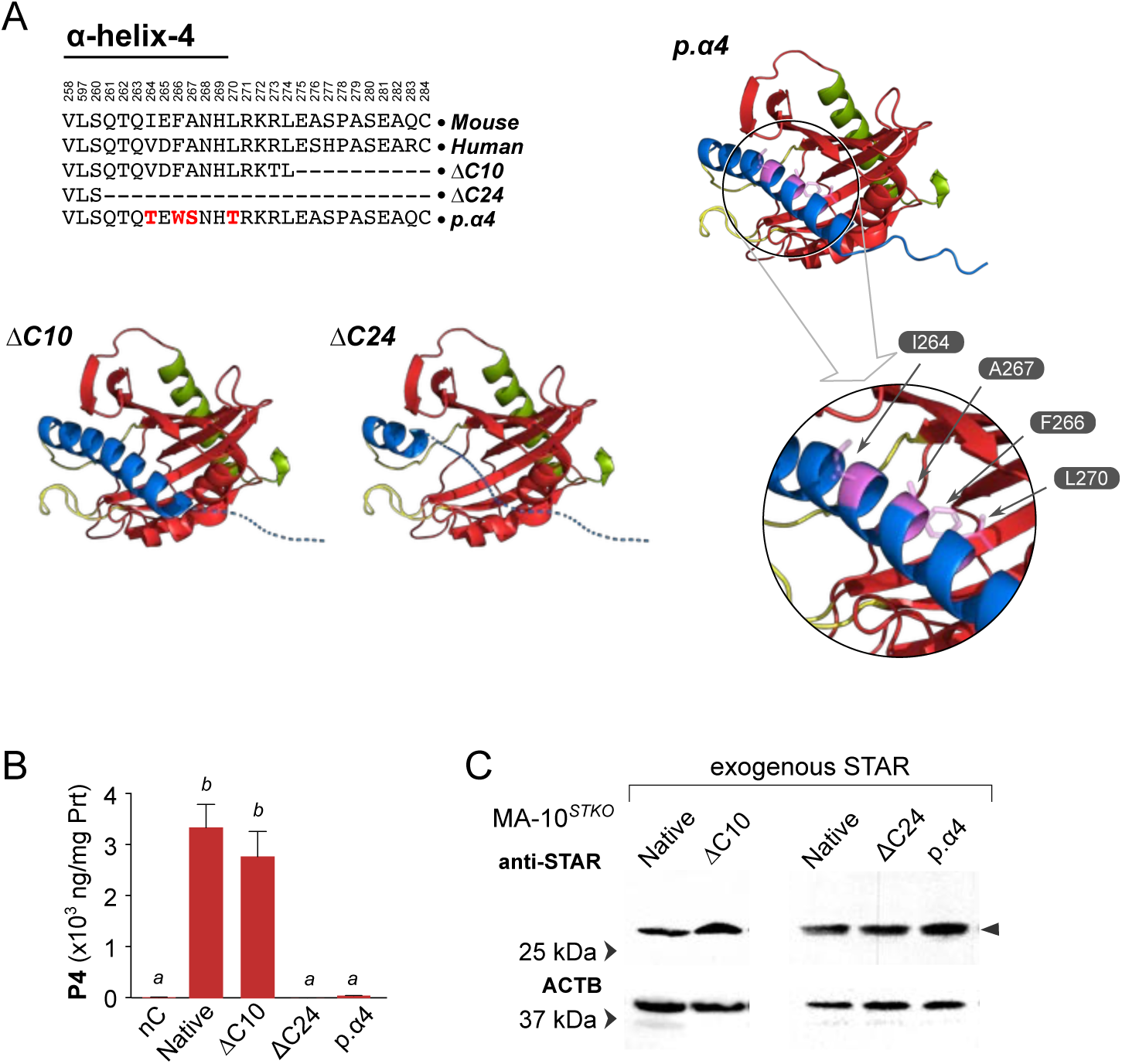
Hydrophobic residues in the C-terminal α-helix 4 of STAR are essential for steroidogenic function. (**A**). The amino acid sequence of the C-terminal α-helix 4 is highly conserved between mouse and human STAR; amino acid numbers are based on the murine sequence. To test its functional importance, we generated three C-terminal mutants: ΔC10, deleting the last 10 amino acids; ΔC24, deleting the last 24 amino acids; p.α4, a substitution mutant in which four key hydrophobic residues—I264, F266, A267, and L270—were replaced with hydrophilic residues (T264, W266, S267, and T270, respectively). Structural diagrams illustrate the extent of C-terminal truncations and locations of targeted hydrophobic residues. **(B)** Functional reconstitution in MA-10^STKO^ cells showed that ΔC10 retained full steroidogenic activity, comparable to native STAR (different alphabets indicate significant difference p<0.05). In contrast, both ΔC24 and p.α4 mutants completely failed to restore progesterone (P4) production, indicating that the integrity of α-helix 4 and its hydrophobic character are critical for function. **(C)** Western blot analysis confirmed comparable expression levels of all STAR variants, ruling out protein instability or differential expression as causes for loss of function.

**Table S1.** Frameshift mutations in STAR-associated lipoid CAH in patients.

|  | **Mutation** | **Case study** |
| --- | --- | --- |
| MTS | Del 7bp5 | https://doi.org/10.1016/j.jpeds.2004.04.052 |
|  | *Del G22_L59 | https://doi.org/10.1210/jc.2012-2865 |
|  | Sub 64-2T>G | https://doi.org/10.1056/NEJM199612193352503 |
|  | Sub 64-1G>T | https://doi.org/10.1210/jc.2003-030345 |
|  | Sub 65-2A>G | https://doi.org/10.1159/000321348 |
| α-helix 1 | Ins A163 | https://doi.org/10.1210/jcem.85.10.6896 |
|  | Ins 3T178 | https://doi.org/10.1210/jcem.82.7.4045 |
|  | Del 201_202CT | https://doi.org/10.1210/jc.2007-1306 |
| α-helix 2 | Ins G246 | https://doi.org/10.1093/hmg/6.4.571 |
|  | Ins G248 | https://doi.org/10.1056/NEJM199612193352503 |
|  | Del T261 | https://doi.org/10.1073/pnas.92.11.4778 |
| β-sheet 6 | Ins 2T549 | https://doi.org/10.1056/NEJM199612193352503 |
|  | Del 13bp564 | https://doi.org/10.1093/hmg/6.4.571 |
| β-sheet 7 | Del 2T593 | https://doi.org/10.1056/NEJM199612193352503 |
| Loop 9 | Ins T643 | https://doi.org/10.1210/jcem.85.10.6896 |
|  | Del C650 | https://doi.org/10.1056/NEJM199612193352503 |
| β-sheet 7 | Ins A677 | https://doi.org/10.1210/jcem.85.10.6896 |
| Loop 11 | Sub 745-1G>C | https://doi.org/10.1177/11795514231167059 |
| Loop 12 | Del A838 | https://doi.org/10.1172/JCI119284 |

\*Mutations directly relevant to this study

**Table S2.** Amino acid substitution mutations in STAR-associated lipoid CAH patients.

|  | **Mutation** | **Case study** |
| --- | --- | --- |
| α-helix 2 | Q77X | https://doi.org/10.1186/s12902-018-0307-6 |
| Loop 5 | R140P | https://doi.org/10.1016/j.jpeds.2004.04.052 |
|  | M144R | https://doi.org/10.1210/jc.2004-1323 |
|  | G146A | https://doi.org/10.1159/000321348 |
|  | W147X | https://doi.org/10.4274/Jcrpe.927 |
|  | N148K | https://doi.org/10.1530/EDM-15-0119 |
| β-sheet 5 | E169G | https://doi.org/10.1056/NEJM199612193352503 |
|  | E169K | https://doi.org/10.1056/NEJM199612193352503 |
|  | E169G | https://doi.org/10.1056/NEJM199612193352503 |
| Loop 7 | *L178Q | https://doi.org/10.3892/mmr.2020.11140 |
|  | R182C | https://doi.org/10.6065/apem.2013.18.1.40 |
|  | R182L | https://doi.org/10.1056/NEJM199612193352503 |
|  | R182L | https://doi.org/10.1056/NEJM199612193352503 |
|  | R182H | https://doi.org/10.1210/jc.2004-1323 |
| β-sheet 6 | V187M | https://doi.org/10.1210/jc.2006-1565 |
|  | R188C | https://doi.org/10.1210/jc.2006-1565 |
| | ΔG189 | https://doi.org/10.1093/hmg/6.4.571 |
|  | R192C | https://doi.org/10.1210/jc.2009-0467 |
|  | R193X | https://doi.org/10.1126/science.7892608 |
| Loop 8 | S195A | http://ci.nii.ac.jp/naid/10007524803 |
| Loop 9 | A203D | https://doi.org/10.1507/endocrj.k05-174 |
|  | Q212X | https://doi.org/10.1093/hmg/6.4.571 |
|  | R217T | https://doi.org/10.1210/jcem.84.11.6118 |
|  | A218V | https://doi.org/10.4274/Jcrpe.927 |
|  | A218V | https://doi.org/10.1056/NEJM199612193352503 |
|  | G221D | https://doi.org/10.1210/jc.2010-0437 |
| β-sheet 8 | M225T | https://doi.org/10.1093/hmg/6.4.571 |
| Loop 11 | D246G | https://doi.org/10.3892/mmr.2020.11140 |
|  | W250X | https://doi.org/10.1210/jcem.84.5.5694 |
| α-helix 4 | Q258X | https://doi.org/10.1126/science.7892608 |
|  | L260P | https://doi.org/10.1210/jc.2005-0874 |
|  | *F267S | https://doi.org/10.1210/jc.2010-0437 |
|  | R272P | https://doi.org/10.4274/Jcrpe.927 |
| | ΔR272 | https://doi.org/10.1056/NEJM199612193352503 |
|  | L275P | https://doi.org/10.1056/NEJM199612193352503 |

\*Mutations directly relevant to this study

**Table S3.**
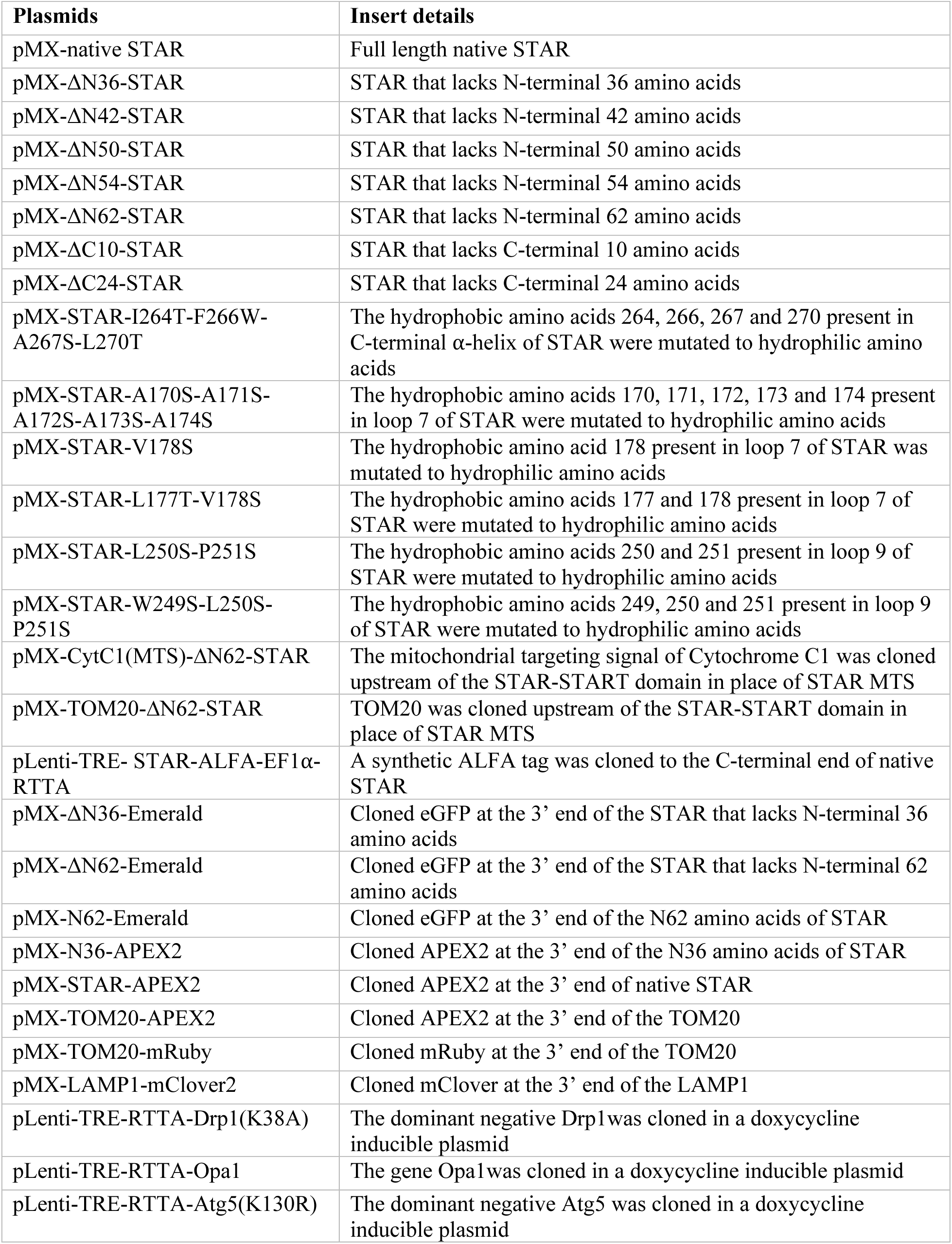
List of the plasmid vectors and transgenes used in this study.

